# Unconventional binding of Calmodulin to CHK2 kinase inhibits catalytic activity

**DOI:** 10.1101/2024.10.08.617309

**Authors:** Christopher R. Horne, Tingting Wang, Samuel N. Young, Toby A. Dite, Hunter G. Nyvall, Sushant Suresh, Katherine A. Davies, Lucy J. Mather, Laura F. Dagley, Gerard Manning, Anthony R. Means, John E. Burke, Janni Petersen, John W. Scott, James M. Murphy

## Abstract

Calmodulin (CaM) serves an essential role in eukaryotic cells as a Ca^2+^ sensor. Ca^2+^ binding leads to conformation changes in CaM that enable engagement of a repertoire of enzymes and the regulation of their catalytic activities. Classically, Ca^2+^-CaM binds to an inhibitory pseudosubstrate sequence C-terminal to the kinase domain in members of the Ca^2+^-CaM dependent protein kinase (CAMK) family, and relieves inhibition to promote catalytic activity. Here, we report an unexpected mechanism by which CaM can bind CHK2 kinase to inhibit its kinase activity. Using biochemical, biophysical, structural mass spectrometry, and cellular approaches, we identify a direct interaction of Ca^2+^-CaM with the CHK2 kinase domain that suppresses CHK2 catalytic activity in vitro and is crucial for cell proliferation in human cells following DNA damage. Our findings add direct suppression of kinase activity to the repertoire of CaM’s functions, complementing the paradigmatic mechanism of promoting kinase activity through autoinhibitory domain sequestration.

## INTRODUCTION

Calmodulin (CaM) is a ubiquitous intracellular protein in eukaryotic cells, which constitutes up to 0.1% of the total proteome and serves a crucial function as a Ca^2+^ sensor^1^. Ca^2+^ is a crucial second messenger in many facets of biology, from muscle contractility to neuronal communication, and its binding to CaM has been likened to a ligand binding to and activating a receptor^1, 2^. CaM binding to four Ca^2+^ ions promotes a conformational change that enables engagement of downstream signaling effectors^3, 4^, including GTPases, GPCRs, adenylyl cyclases, pseudokinases, phosphatases, lipid and protein kinases^5^, to modulate their activities. The cellular functions of these target proteins can be regulated by CaM via a range of mechanisms, such as by CaM dictating their localization within cells, the occlusion of substrate or partner protein binding, or by modulating a binding partner’s catalytic activity.

CaM is composed of two globular Ca^2+^ binding domains separated by a solvent exposed central helix^1^. Each globular domain contains two EF-hand Ca^2+^ binding motifs that bind Ca^2+^ with low micromolar affinity. The CaM binding sequences (CaMBS) of the target proteins each contain 15-30 amino acids^3, 4^ and bind CaM with affinities ranging from low nanomolar to low micromolar^6, 7^. While the precise target sequence is highly variable between among the plethora of interactors, there are two common principles guiding CaM binding. Firstly, the CaMBS contains a cluster of 2-3 basic residues, which defines the polarity of the binding of the sequence to CaM. Secondly, the CaMBS typically forms a helix and projects hydrophobic anchor residues into non-polar clefts within CaM (reviewed in ref.^1, 5, 8^). Structural studies have revealed enormous diversity in the mode of CaMBS binding to CaM, where the periodicity of the helix allows the anchor residues to occur in diverse positions. Accordingly, anchor residue positions of 1; 4 or 5; 8 or 10; 13, 14, 16 or 17 have been identified in a variety of CaMBS, often with anchor residues contributed beyond the helix^5^. Our current understanding allows for predictions of conventional CaMBS, although prediction of non-canonical CaMBS remains elusive and relies on experimental determination.

Ca^2+^-CaM serves a crucial function in the activation of the Ca^2+^-CaM-dependent protein kinase (CAMK) family following Ca^2+^ influx into cells^2^. The CAMK family is a large and diverse kinase subfamily, which totals 74 family members^9^. Among these, the activation of myosin light chain kinase (MLCK), and CAMK1 (including CaMKI and CaMKIV kinases) and CAMK2 (including CaMKII kinases) families are known to occur via a two-step Ca^2+^-CaM-dependent process, although if and how Ca^2+^-CaM activates other CAMK kinases is poorly understood. In the canonical mechanism, Ca^2+^-CaM binds to a sequence C-terminal to the kinase domains, which overlaps an autoinhibitory sequence that occludes the active site. Ca^2+^-CaM binding displaces the autoinhibitory sequence from the active site, thereby activating the kinase. The mechanism is best described for CaMKII^10^, which along with DAPK1^11^, DAPK2^12^ and GRK5^13^, are the only four structures of Ca^2+^-CaM bound kinase structures reported thus far. The CaMKII autoinhibitory sequence binds in a helical conformation to the kinase domain C-lobe to prevent CaMKII activation by trans-autophosphorylation. Ca^2+^-CaM binds to a sequence overlapping the C-terminal autoinhibitory sequence, leading to unfolding of the N-terminus of the autoinhibitory helix and formation of an extended helix that then binds CaM^10^. CaM binding thus induces dissociation of the autoinhibitory sequence from the CaMKII kinase domain C-lobe, which in turn allows activation via autophosphorylation to ensue^14^. Similarly, in the case of CAMK1 family kinases, their upstream activating kinases, the CaMKKs, are able to access and phosphorylate the CaMK activation loops to activate their kinase activity when the autoinhibitory sequence is sequestered by Ca^2+^-CaM^2^. That Ca^2+^-CaM activates both CAMK and CAMKK kinases is intriguing, and indicates a crucial function for Ca^2+^-CaM in signal fidelity and amplification upon pathway engagement. Based on the limited structural data and extensive biochemical studies, the CaM binding sequence C-terminal to the kinase domain is the target of Ca^2+^-CaM in both the CaMK and CaMKK kinases, and has served as a paradigm for how CaM can promote kinase catalytic activity for decades. However, whether others among the vast array of CAMK kinases, and kinases from other families, are Ca^2+^-CaM regulated has remained unclear to date.

A member of the CAMK group – Checkpoint kinase 2 (CHK2 or CHEK2) – is a highly-studied protein kinase owing to its essential role in regulating cell cycle progression following DNA damage^15^. CHK2 is a multidomain protein comprising an N-terminal SQ/TQ cluster domain (SCD), a Forkhead-associated (FHA) domain, and a C-terminal Ser/Thr kinase domain. Following DNA damage, the upstream kinase, ATM, phosphorylates Thr68 within the CHK2 SCD, which introduces a site for CHK2 FHA domain binding^16, 17^. CHK2 is reported to form a transient homodimer, presumably mediated by FHA domain binding to pT68 *in trans*^18^, which in turn leads to CHK2 activation loop phosphorylation on T383 and elevated kinase activity^19^. CHK2 does not harbour a canonical autoinhibitory Ca^2+^-CaM binding sequence yet was previously proposed to be activated by Ca^2+^-CaM^20^, although these data were far from conclusive. Here, we identify an unexpected role for Ca^2+^-CaM as an inhibitor of CHK2 kinase activity that functions via direct engagement of the CHK2 kinase domain. Structural mass spectrometry, modelling, and mutational studies pinpoint a bidentate interaction of the two EF hand domains of CaM with the N- and C-lobes of CHK2, centered on the catalytic loop. Moreover, disruption of the CaM binding interface via CRISPR mutation of the critical residue in CHK2, K373, impeded cell proliferation and ablated proliferation following DNA damage in human cells. This functionally-crucial lysine is highly conserved amongst orthologs, which suggests an ancestral function for CaM as the sensor of Ca^2+^ flux that attenuates CHK2 activity and cell proliferation. These findings expand the repertoire of mechanisms by which CaM can regulate the catalytic activity of protein kinases, raising the prospect that CaM may function allosterically to modulate the activities of enzyme targets more broadly in nature.

## RESULTS

### Ca^2+^-CaM directly binds and inhibits CHK2 catalytic activity

Upon Ca^2+^ binding, CaM typically binds to an autoinhibitory site C-terminal to the kinase domain in CAMK family kinases to promote catalytic activity (**Fig. 1A**). A control CAMK family kinase, CaMKK2, which is reported to contain a conventional CaMBS that is autoinhibitory^21^ (**Fig. 1A**), bound CaM in a Ca^2+^-dependent manner by Far Western assay (**Fig. 1B**). Surprisingly, like CaMKK2, immobilized CHK2 similarly bound CaM in a Ca^2+^-dependent manner that was compromised by the Ca^2+^ chelator, EGTA (**Fig. 1B**). CHK2 lacks a predicted CaMBS, indicating that CaM engages CHK2 via an unconventional binding mode. We tested the impact on CHK2 catalytic activity by performing radiometric kinase assays on full-length recombinant CHK2 **(Fig. 1C**), revealing that CHK2 activity was attenuated nearly two-fold by CaM. Comparable results were obtained using an orthogonal fluorometric assay, providing further support for CaM interacting with CHK2, and inhibiting its catalytic activity (**Supp. Fig. 1A**). We next asked whether CHK2 and CaM directly interact using surface plasmon resonance (SPR). We immobilized full-length recombinant CHK2 to the sensor surface via a fused, biotinylated AviTag motif and examined binding to Ca^2+^-CaM as the analyte. These experiments validated a direct interaction between Ca^2+^-CaM and CHK2, with a low micromolar affinity (K_d_ = 21.6 ± 4.7 μM) (**Supp. Fig. 2**).

**Figure 1.**
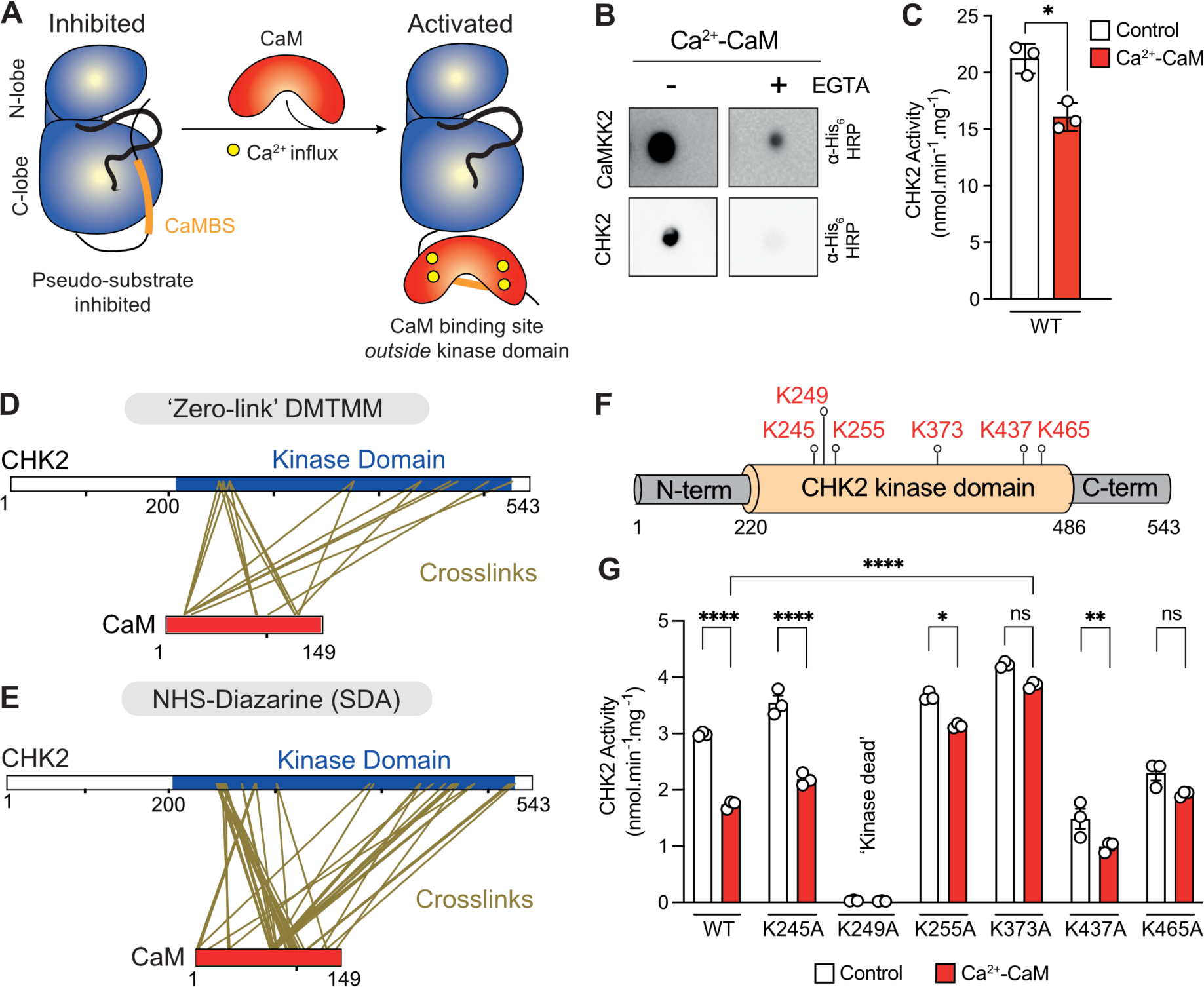
Ca^2+^-CaM interacts with human CHK2 to inhibit kinase activity via interaction with the kinase domain. **(A)** Schematic of the conventional mode of Ca^2+^-CaM activation in members of the Ca^2+^-CaM-dependent protein kinase (CaMK) family. Upon Ca^2+^ influx, Ca^2+^-CaM binds a linear sequence (CaMBS; orange) that is proximal to the CaMK kinase domain (blue). This interaction restores activity by sequestering an autoinhibitory pseudosubstrate sequence, which formerly obstructed the active site. **(B)** Far-Western blots show that recombinant CHK2 and control kinase, CaMKK2, interact with His_6_-CaM in a Ca^2+^-dependent manner (Ca^2+^: 500 μM), as binding is abolished in the presence of the Ca^2+^-chelator, EGTA (10 mM). The interaction was probed using anti-His_6_ HRP antibody. Data are representative of two independent replicates. **(C)** CHK2 is inhibited by Ca^2+^-CaM. CHK2 was immunoprecipitated and activity was measured by radiometric assay in the presence or absence of 100 μM CaCl_2_, 1 μM CaM and 200 μM [^32^P]-γ-ATP for 10 min. Data represent mean ± SD; n = 3. Statistical analysis was performed by paired t-test; * signifies P<0.1. **(D)** Schematic of the amino acid residues of full-length CHK2 and CaM (red) chemically cross-linked by DMTMM (0 Å spacer) and **(E)** photoactivatable crosslinker, NHS-Diazarine (SDA; 3.9 Å spacer), following mass-spectrometry analysis. Cross-links (gold) are only observed within the CHK2 kinase domain (blue) for both chemical crosslinkers. **(F)** Six lysine residues in CHK2, identified by DMTMM chemical crosslinking are mapped to the domain architecture of CHK2. **(G)** Ca^2+^-CaM inhibition of wild-type CHK2 kinase domain (210-531) and cross-linked lysine mutants. CHK2 kinase domain was recombinantly expressed and activity was measured in the presence or absence of 100 μM CaCl_2_, 1 μM CaM and 200 μM [^32^P]-γ-ATP for 10 min. Data represent mean ± SD; n = 3. Statistical analysis was performed by two-way ANOVA; *, ** and **** signifies P<0.1, P<0.01 and P<0.0001, respectively; n.s = non-significant.

Having established a direct inhibitory interaction between Ca^2+^-CaM and CHK2, we mapped the site of interaction using crosslinking mass spectrometry with the zero-length crosslinker, DMTMM (**Fig. 1D**), and the photoactivatable crosslinker, SDA (**Fig. 1E**). Using either crosslinker, interactions with CaM were confined to the CHK2 kinase domain, even though full length recombinant CHK2 was used in these studies. DMTMM crosslinking identified 6 lysines in CHK2 that were proximal to CaM in the complex (**Fig. 1F**). We examined the role of these CHK2 lysines in Ca^2+^-CaM binding by preparing recombinant CHK2 kinase domain (210-531) containing individual Lys to Ala substitutions (**Supp. Fig. 1B**) to examine their susceptibility to Ca^2+^-CaM inhibition (**Fig. 1G**). One substitution, K249A, mutated the canonical ATP-binding lysine of the VAIK motif, which eliminated its catalytic activity, as expected. K245A CHK2 retained sensitivity to Ca^2+^-CaM inhibition, whereas K255A, K373A, K437A and K465A CHK2 exhibited diminished sensitivity to Ca^2+^-CaM inhibition. Importantly, introduction of the K255A and K373A CHK2 mutations did not compromise basal catalytic activity, while K465A caused a modest attenuation. Retention of basal CHK2 catalytic activity, but loss of Ca^2+^-CaM modulation, supports roles for these residues in Ca^2+^-CaM engagement, rather than in CHK2 catalysis. To ensure these differences in Ca^2+^-CaM sensitivity were not attributable to compromised protein folding, we evaluated each variant’s thermal stability by differential scanning fluorimetry. All mutant CHK2 (210-531) constructs, except K249A and K465A CHK2, exhibited melting temperature (*T*_m_) that were comparable to wild-type (ranging from 39 to 44 °C; **Supp. Fig. 1C**), demonstrating that these constructs were correctly folded. K465A displayed a lower *T*_m_ (28 °C) suggesting partial destabilization or increased flexibility of this mutant, which may explain the lower basal catalytic activity (**Fig. 1G**). An apparent *T*_m_ could not be obtained for K249A CHK2 due to high background fluorescence. Reduced thermal stability for K249A CHK2 is not surprising, because this mutation would disrupt interaction of the ATP-binding Lys with the αC-helix Glu (E274), an interaction typical of a canonical kinase active state.

### CaM wraps around the CHK2 kinase domain to inhibit activity

We next detailed the interaction of the CHK2 kinase domain with Ca^2+^-CaM using biophysical methods. Hydrogen-deuterium exchange mass spectrometry (HDX-MS) is a technique that measures the exchange rate of amide hydrogens, which acts as a readout of protein secondary structure dynamics^22^. HDX-MS can be employed to identify both direct and allosteric conformational changes accompanying ligand binding. HDX-MS has been used extensively to identify conformational changes accompanying protein kinase regulation, inhibition and activation^23, 24, 25^. We used HDX-MS to map regions within full length recombinant CHK2 with altered hydrogen-deuterium exchange in the presence of Ca^2+^-CaM, identifying regions centered on, and flanking, the active site cleft (**Fig. 2A, B**). Deuterium exchange was decreased in peptides encompassing residues 72-96, 97-106 and 123-130 in the FHA domain, in addition to peptides 203-218 and 205-220 at the junction with the kinase domain, and 353-363, 437-452 and 439-452 in the kinase domain (**Supp. Fig. 3**). These decreases in exchange are consistent with the crosslinking mass spectrometry data (**Fig. 1D, E; 2C**), where CaM wrapping around the CHK2 kinase domain would be expected to induce changes in hydrogen-deuterium exchange in the catalytic loop and flanking regions of the kinase domain. Accordingly, in the presence of Ca^2+^-CaM, reduced HDX was observed in peptides at the flexible FHA-kinase domain junction N-terminal to the kinase domain and at the neighboring region in the FHA domain, and in the αG and αG′ helices at the foot of the kinase domain C-lobe (**Fig. 2B-C**). Mapping the electrostatic potential to the molecular surface of the CHK2 kinase domain reveals extensive patches of positive charge that we predicted to bind CaM from our crosslinking experiments, which complement the highly-negatively-charged surface of CaM (**Fig. 2D**). Our crosslinking mass spectrometry and HDX-MS data were supported by an AlphaFold2 model of the CaM:CHK2 kinase domain complex, in which a bidentate interaction of CaM with the N- and C-lobes of CHK2, centered on the CHK2 catalytic loop, was observed (**Fig. 2E**).

**Figure 2.**
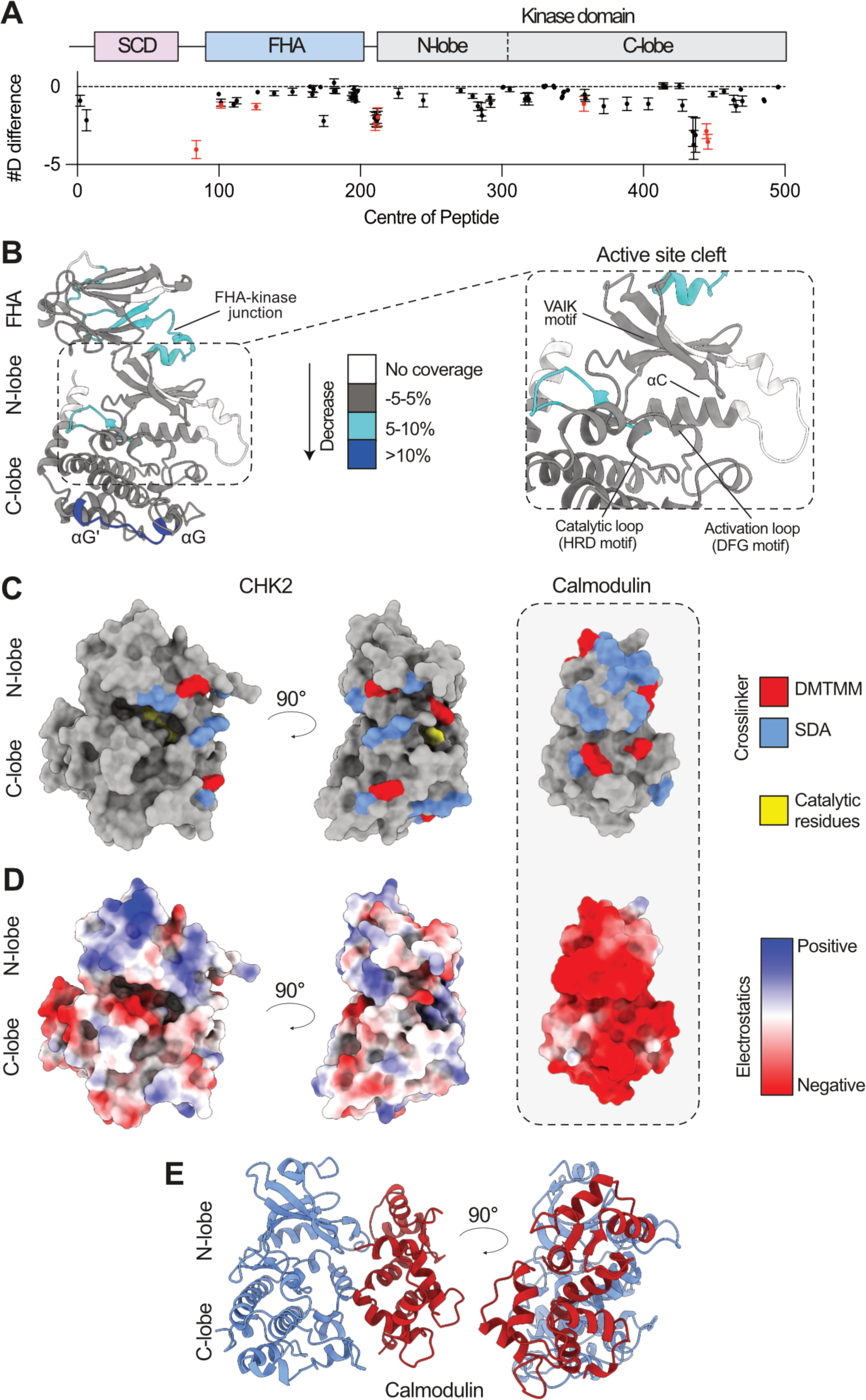
Mapping the Ca^2+^-CaM binding surface of human CHK2. **(A)** Quantitative analysis of deuterium exchange differences of human CHK2 in the presence of Ca^2+^-CaM, following hydrogen-deuterium exchange mass spectrometry (HDX-MS). The graph shows the #D difference in deuterium incorporation for CHK2 over the entire time course. Each point represents an individual peptide, with those colored in red showing a significant change (greater than 0.40 Da and 5% difference at any timepoint, with a two tailed t-test; p<0.01, n=3), with error bars showing standard deviation. CHK2 domain architecture is annotated above. **(B)** Significant differences in deuterium exchange are mapped onto an AlphaFold model of CHK2 and are color-coded according to the legend. Zoomed view of the CHK2 active site cleft is shown on the right. **(C)** Crosslinking mass spectrometry data is mapped onto an AlphaFold model of CaM:CHK2. ‘Zero-link’ DMTMM and SDA crosslinks are shown in red and blue, respectively. Key catalytic residues of CHK2 are shown in yellow. **(D)** The electrostatic potential mapped onto the molecular surface of CHK2 shows a cluster of positive charge that align with residues in (C) and complement the negatively-charged surface of CaM. **(E)** AlphaFold model of human CHK2 kinase domain (blue) in complex with CaM (red).

### CaM binding K373 in CHK2 is crucial to cellular function

We next sought to establish whether the key CaM-binding residues in CHK2 identified by crosslinking mass spectrometry, and validated as important for Ca^2+^-CaM interaction biochemically, are crucial to CHK2’s cellular function. CHK2 is highly evolutionarily-conserved, with orthologs known in animals and fungi (**Fig. 3A**). Examination of sequences identified K373 of human CHK2 as almost universally conserved, with the only deviation being *Trichoplax adhaerens* where the Lys was conservatively substituted with Arg (**Fig. 3A**). The extensive conservation of K373 suggested a conserved functional role, which warranted further investigation in a cellular context. By contrast, the other human CHK2 residues implicated in interaction with CaM, K255 and K465 (**Fig. 1D, F, G**), are highly conserved as Lys or Arg in mammalian CHK2 orthologs, but are poorly conserved in other eukaryotes, and so less likely to serve an ancestral role in CaM recognition than K373. Accordingly, we focused our cellular examination of CaM regulation of CHK2 on the K373A mutant. Using CRISPR, we introduced an Ala substitution for K373 into the diploid, hTERT-immortalized, retinal pigment epithelial (RPE) cells^26^ (**Fig. 3B; Supp.** Fig. 4), which are routinely used to study the cell cycle^27^. Following CRISPR-knockin, analysis of cell proliferation using IncuCyte live cell imaging revealed that, while *CHK2^K373A^* RPE cells grew, they exhibited muted proliferation compared to wild-type RPE cells (**Fig 3C, D**). These growth deficiencies for CHK2 were accentuated by exposure to the DNA damaging agent, Camptothecin (CPT), a topoisomerase I inhibitor^28^ (**Fig. 3E**). Cell proliferation of *CHK2^K373A^* RPE cells was significantly reduced in the presence of 10nM and 100nM CPT compared to wild-type RPE cells (**Fig. 3F**), as was evident in end-point micrographs (**Fig. 3G**). These data implicate the CHK2 activation loop – centred on K373 – as serving an essential role in CaM engagement, where loss of CaM interaction compromises the DNA damage response and cell cycle progression in response to Ca^2+^ flux.

**Figure 3.**
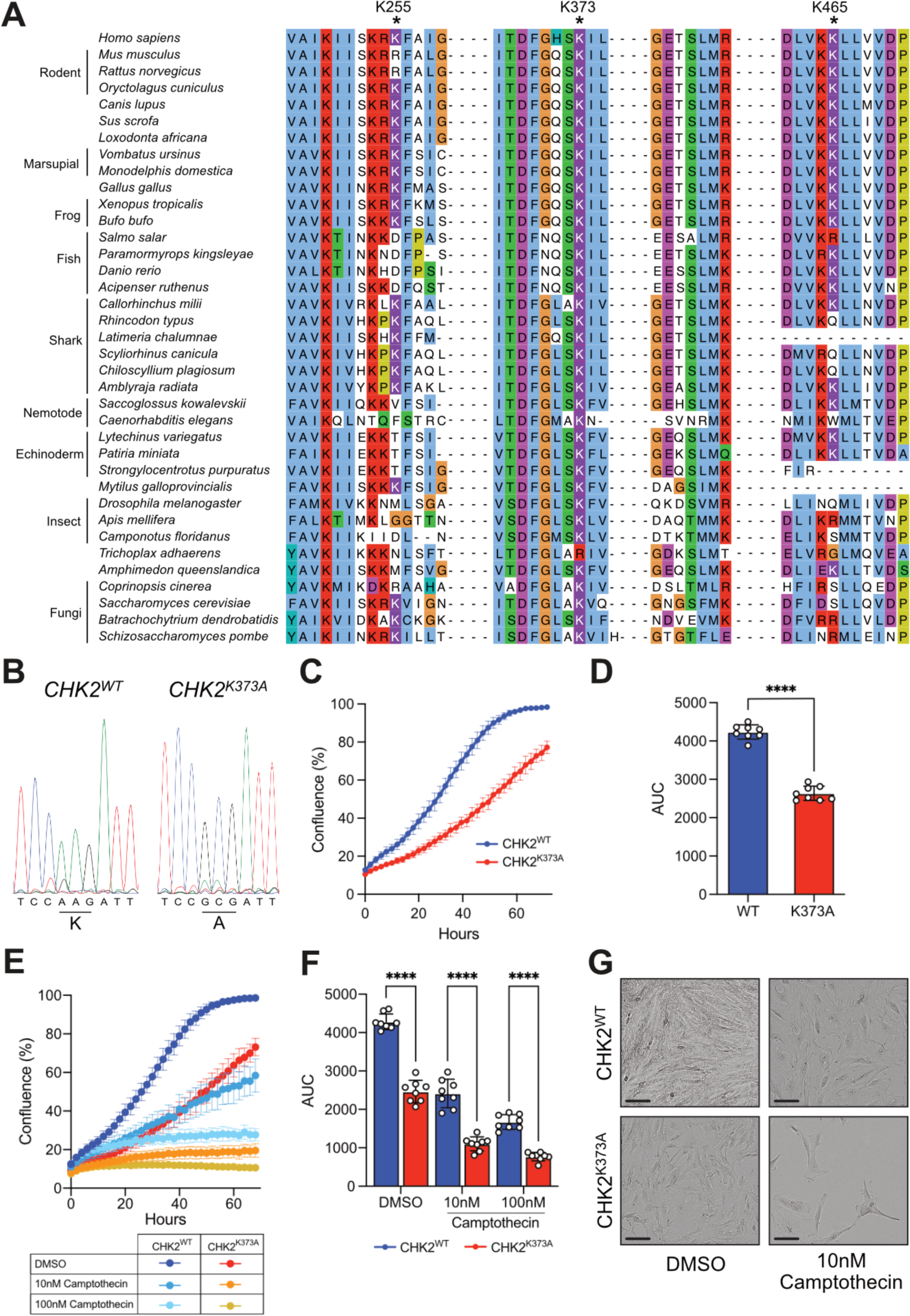
Mutation of evolutionarily-conserved K373 of human CHK2 compromises cell proliferation. **(A)** Multiple sequence alignment of CHK2 orthologs, which highlights the conservation of CHK2 residues in animals and fungi (K255, K373 and K465; asterisk) implicated in interaction with Ca^2+^-CaM. Residues in the alignment are colored according to the Clustal coloring scheme, which highlights chemical nature and patterns of conservation. **(B)** Chromatograms from Sanger sequencing confirm the successful integration of nucleotide changes for K373A in hTERT-immortalized RPE cells by CRISPR/Cas9. **(C)** Real-time proliferation analysis of wild-type and CRISPR-edited RPE cells using IncuCyte live-cell imaging, and **(D)** corresponding area under the curve. Data represent mean ± SD; n = 8. Statistical analysis was performed by paired t-test; **** signifies P<0.000. **(E)** Real-time proliferation analysis of wild-type and CRISPR-edited RPE cells cultured in the presence of DNA damaging agent, Camptothecin (CPT), at 10nM and 100nM, and **(F)** corresponding area under the curve. Data represent mean ± SD; n = 8. Statistical analysis was performed by two-way ANOVA; **** signifies P<0.0001. **(G)** Raw micrographs of cell confluence exported from the IncuCyte SX3 system after 72 h incubation; the scale bars (black) represent 100 µm.

## DISCUSSION

Kinases of the CAMK1 and CAMK2 families are known to be activated by Ca^2+^-CaM sequestering an autoinhibitory sequence C-terminal to the kinase domain, to relieve autoinhibition of the kinase domain (**Fig. 4**). Elsewhere among the CAMK family, few members are predicted to harbor CaMBS and, rather than association with Ca^2+^-CaM regulation, have been classified in the CAMK family owing to their sequence homology. Accordingly, in the absence of CaMBS, whether they are CaM regulated has not been established and, if they are, the underlying mechanism remains undetailed.

**Figure 4.**
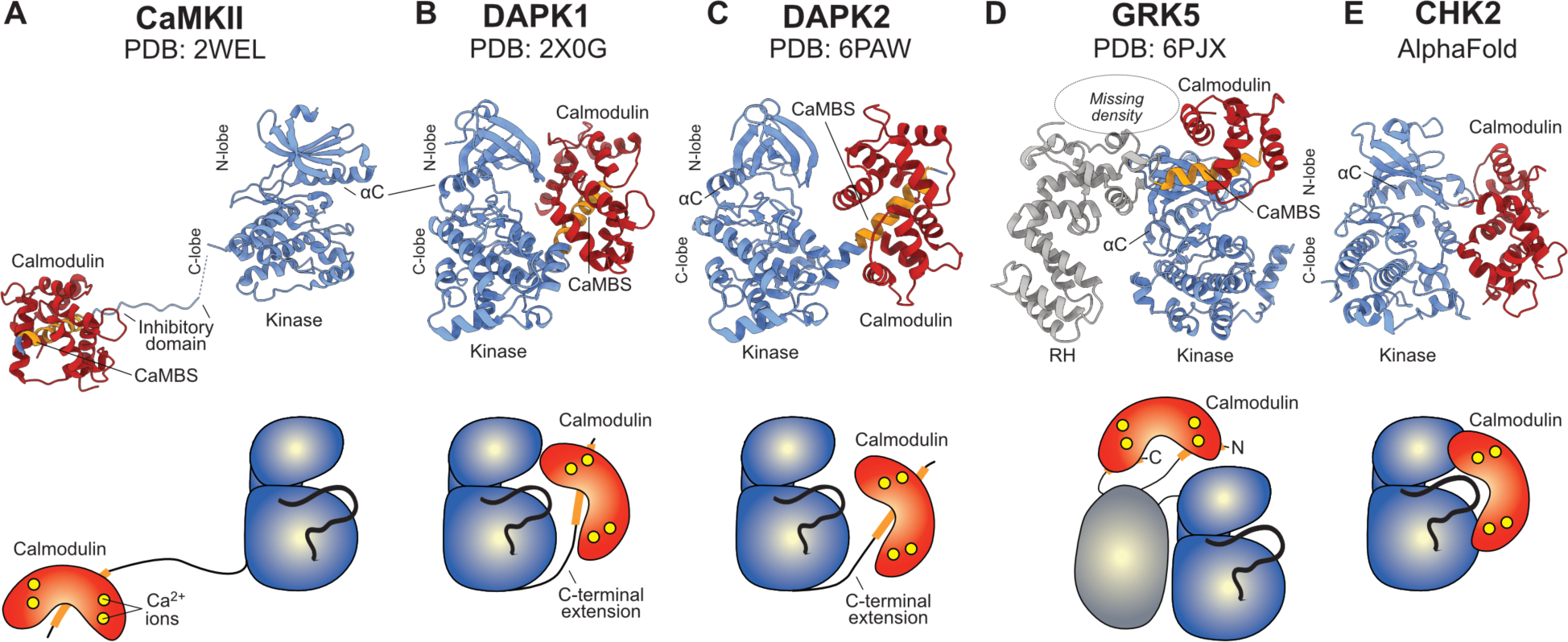
Comparison of known modes of Ca^2+^-CaM regulation of protein kinases. **(A)** Ca^2+^-CaM-dependent protein kinase II (PDB 2WEL)^10^; **(B)** Death-associated protein kinases (DAPK)-1 (PDB 2X0G)^11^; and **(C)** DAPK2 (PDB 6PAW)^12^ harbor a conventional CaMBS C-terminal to the protein kinase domain. (D) GPCR kinase 5 (GRK5; PDB 6PJ6)^13^ interacts with Ca^2+^-CaM via two helical regions N- and C-terminal to the protein kinase domain. (E) Ca^2+^-CaM directly interacts with the protein kinase domain of CHK2 (AlphaFold model; from this study). Each mode of Ca^2+^-CaM regulation is illustrated by cartoon ribbon (top panel) and schematic representation (lower panel). The protein kinase domain is colored blue, while other domains are grey. CaM and CaMBS are colored red and orange, respectively. Ca^2+^ ions are colored yellow.

Here, we identify binding of Ca^2+^-CaM to the kinase domain of CHK2 (**Fig. 4**), which, unlike typical Ca^2+^-CaM activation of CAMK family kinases^2, 14^, inhibits CHK2 activity. There are precedents for direct engagement of Ca^2+^-CaM with the kinase domain or sites N-terminal to the kinase domain from the world of plant kinases, which promote catalytic activity, although it is important to note that these attributions are based primarily on Ca^2+^-CaM binding to synthetic peptides rather than native kinase domains^29, 30, 31^. Here, we identified the catalytic loop and flanking regions in the N- and C-lobes of the CHK2 kinase domain as the site of CaM binding. K373 in the CHK2 kinase domain activation loop, which is almost absolutely conserved throughout evolution, was essential for cell cycle progression following DNA damage. CHK2 is known to phosphorylate CDC25C, BRCA1, HDMX, PLK1 and TP53^32, 33, 34, 35^, which antagonizes their promotion of cell cycle arrest and DNA repair. Accordingly, blockade of CHK2-mediated effector phosphorylation will relieve their inhibition and enable cell cycle progression and DNA repair (**Fig. 5**). Importantly, our binding experiments identified a K_d_ ∼20μM for interaction of Ca^2+^-CaM to CHK2. Such an affinity would allow lability, such as during flux in Ca^2+^ levels within the nucleus that would modulate CaM conformation towards the CHK2 binding-competent form, as well as ensure inhibitory CaM binding only upon high local concentrations in the nucleus. The best characterized mode of attenuation of the activities of CHK2 and other CAMK kinases is phosphatase-mediated dephosphorylation to disarm their kinase activity or transactivation^36, 37^. Here, we add direct binding to the kinase domain as another mechanism by which Ca^2+^-CaM can negate CHK2 activity and block phosphorylation of downstream substrates to enable cell cycle progression and DNA repair.

**Figure 5.**
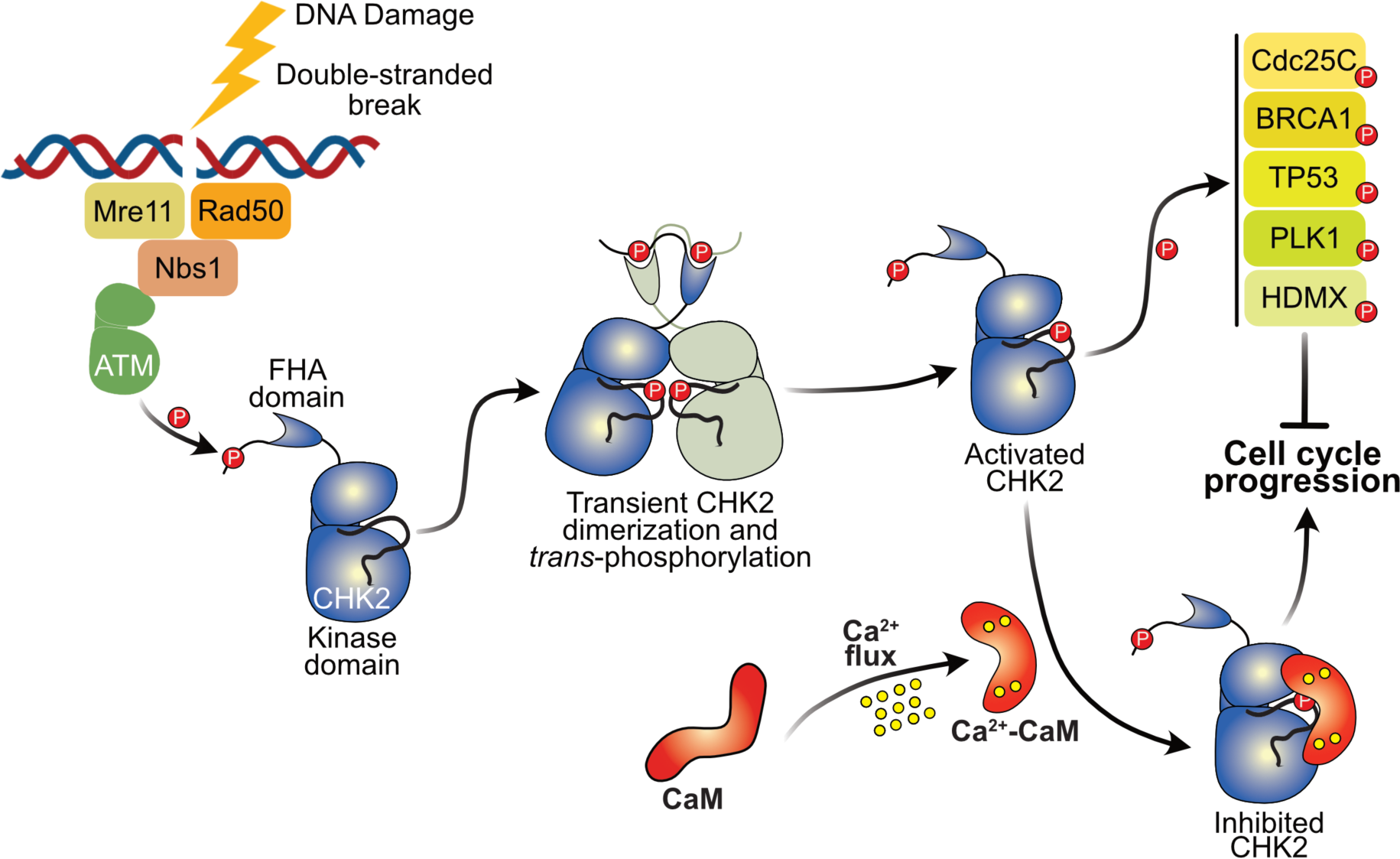
Schematic model of CHK2 activation and CaM-mediated regulation of cell cycle progression. Following DNA damage, the upstream kinase, ATM (green), phosphorylates Thr68 within the CHK2 SCD, which leads to the formation of a transient homodimer, mediated by the CHK2 FHA domains binding to pThr68. This dimerization results in CHK2 activation loop phosphorylation on T383 *in trans* and increased catalytic activity. Following activation, CHK2 phosphorylates multiple substrates, including CDC25C, BRCA1, HDMX, PLK1 and TP53, to promote cell cycle arrest and impede DNA repair. Upon sensing increased Ca^2+^ levels, Ca^2+^-CaM can directly interact with the CHK2 kinase domain to inhibit catalytic activity and enable cell cycle progression and DNA repair.

Broadly, our findings identify an unprecedented mechanism by which Ca^2+^-CaM can regulate kinase activity. Typically, Ca^2+^ laden CaM wraps around helical peptide sequences with hydrophobic residues in the target sequence inserting into an anchor pocket in addition to one or more regularly spaced hydrophobic pockets in CaM^5^. The helix connecting the two EF-hand domains of CaM allows for some flexibility in binding mode, such that the spacing of hydrophobic residues in the target sequence can vary by one or more helical turns, and dictate whether Ca^2+^-CaM binds in a compact or elongated form. Structural knowledge of these binding modes has enabled prediction of sequences that conform to typical CaMBS’s. However, knowledge of whether Ca^2+^-CaM binds three-dimensional surfaces, rather than helical motifs, is very limited. Even in the case of the CAMK family kinase, DAPK1, Ca^2+^-CaM binds principally to the autoregulatory helix C-terminal to the kinase domain, and this complex contacts the kinase domain^11^. Hydrogen bonds between the N-terminal EF-hand and the N-lobe of the DAPK1 kinase domain leads to a head-to-head arrangement of the kinase domain and Ca^2+^-CaM. Superficially the CaM:DAPK1 assembly resembles our model for Ca^2+^-CaM binding to the CHK2 kinase domain, although Ca^2+^-CaM binding to DAPK1 promotes catalytic activity by preventing autoinhibitory sequence binding to the DAPK1 substrate binding site and Ca^2+^-CaM does not occlude substrate binding.

While precise mapping of the CaM:CHK2 interface awaits a high resolution structure, our data suggest Ca^2+^-CaM would inhibit CHK2 activity by occluding the substrate binding site because Ca^2+^-CaM binding is centred on the CHK2 catalytic loop. A high resolution structure will likely be important to defining whether Ca^2+^-CaM imposes more severe structural perturbations, such as by freezing CHK2 in an inactive conformation or disrupting the regulatory “R”-spine of hydrophobic interactions required for an active conformation^38^. Nevertheless, our data support the idea that Ca^2+^-CaM binding to CHK2 acts downstream of CHK2’s activation through homodimerization rather than acting on homodimers themselves. CHK2 dimerization mediated by the N-terminal FHA domain binding *in trans* is reported to be a key step in activation, however Ca^2+^-CaM inhibited activation of autophosphorylated CHK2. Indeed, even at high CHK2 and Ca^2+^-CaM concentrations for size-exclusion chromatography in purification, the elution volume corresponded to 1 CHK2: 1 CaM, rather than the binding of Ca^2+^-CaM to a CHK2 dimer. Collectively, our data indicate the Ca^2+^-CaM acts on monomeric CHK2 to block its activity towards substrates.

Our findings highlight an underappreciated role for Ca^2+^-CaM as an inhibitor of protein kinase activity, which complements its well-established role as an activator of CAMK kinase activities via sequestration of autoinhibitory sequences. Additionally, it remains of interest to understand whether kinases, like CHK2 in the CAMK family and beyond, which lack a predicted CaMBS, can be regulated by Ca^2+^-CaM binding through non-canonical binding modes, such as the mode we describe here: direct kinase domain binding. Whether these additional modes of kinase regulation by Ca^2+^-CaM serve ancestral functions remains of outstanding interest for future studies. Our findings underscore the structural and functional plasticity of Ca^2+^-CaM as a regulator of enzymatic activities and suggest that the diversity of Ca^2+^-CaM’s allosteric activities may be greater than currently documented.

## MATERIALS AND METHODS

### Expression constructs

The gene coding for full-length human CHK2 (Uniprot O96017) was synthesised by Gene Universal (Delaware, USA) and sub-cloned into the mammalian expression vector, pcDNA3.1(-), using the restriction sites XhoI and HindIII with an N-terminal FLAG tag. Individual Ala substitutions were also synthesised (Gene Universal) and subcloned into pcDNA3.1(-) as XhoI-HindIII fragments with an N-terminal FLAG tag.. Full-length CHK2 and the CHK2 kinase domain (residues 210-531) were sub-cloned into the bacterial expression vector, pGEX-2-TEV^39^, as an in-frame fusion with a TEV protease-cleavable N-terminal GST tag using InFusion cloning (Takara). For SPR studies, full length CHK2 was subcloned into the insect expression vector, pFastBac GST, as an in-frame fusion with a TEV protease-cleavable N-terminal GST tag and C-terminal AviTag using InFusion cloning. The gene coding for full-length CaM (Uniprot P0DP23) was synthesised by IDT (Iowa, USA) as a gBlock and sub-cloned into the bacterial expression vector, pPROEX Htb (Life Technologies), as an in-frame fusion with a TEV protease-cleavable N-terminal hexahistidine tag. Oligonucleotide sequences are presented in **Supp. Table 1**. Insert sequences were verified by Sanger sequencing (AGRF, VIC, Australia).

### Transient Expression and Immunoprecipitation

For initial binding assays, full-length CHK2 (wild-type) was expressed in HEK293T cells grown in DMEM (Thermo Fisher) media, supplemented with 8% (v/v) Fetal Calf Serum (FCS; Sigma). At 37 °C with 5% CO_2_. The cells were transfected at 60% confluency using FuGene HD (Roche Applied Science) with 2 µg of pcDNA3.1(-) plasmid containing C-terminal FLAG-tagged human CHK2. After 48 h, transfected cells were harvested by rinsing with ice-cold PBS, followed by rapid lysis *in situ* using lysis buffer (50 mM Tris-HCl pH 7.4, 1% (v/v) Triton X-100, 1 mM PMSF, 1 mM EDTA, 150 mM NaCl, 2 mM Sodium Vanadate, 10 mM NaF) containing a Complete protease inhibitor tablet (Roche). Insoluble debris were removed by centrifugation and supernatants were mixed with 100 μL of anti-FLAG M2 agarose (50% v/v; Millipore) pre-equilibrated in lysis buffer, followed by successive washes in lysis buffer containing 1 mM NaCl, and finally resuspended in 50 mM HEPES, pH 7.4. CHK2 was eluted off the beads by incubating overnight at 4 °C with 100 µL of FLAG peptide (1 mg/mL) in 50 mM HEPES, pH 7.4. Total protein content was quantified using the BCA Protein Assay (Thermo Fisher Scientific), according to manufacturer’s instructions.

### Recombinant Expression and Purification

Full-length CHK2 (residues 2-531), CHK2 kinase domain (residues 210-531) and CaM were primarily expressed in *E. coli* BL21-CodonPlus-RIL (Agilent) cells cultured in Super Broth supplemented with ampicillin (100 µg/mL) at 37 °C with shaking at 220 rpm to an OD_600_ of ∼0.6–0.8. Protein expression was induced by the addition of isopropyl β-D-1-thiogalactopyranoside (IPTG; 250 µM) and the temperature was lowered to 18 °C for incubation overnight.

CHK2 cell pellets were resuspended in GST buffer (20 mM HEPES pH 7.5, 200 mM NaCl, 10% v/v glycerol), supplemented with cOmplete Protease Inhibitor (Roche) and 0.5 mM Bond-Breaker TCEP [tris-(2-carboxyethyl)phosphine] (ThermoFisher Scientific) and were lysed by sonication, before the lysate was clarified by centrifugation (40,000 × *g*, 45 min, 4 °C). The proteins were maintained at 4 °C throughout purification. The clarified lysate was incubated with Glutathione Xpure Agarose resin (UBP-Bio) pre-equilibrated in GST buffer for >1 h on rollers at 4 °C, before the beads were pelleted at 500 × *g* and washed extensively with GST buffer. CHK2 was further purified by cleaving the GST tag (on resin) with recombinant His_6_-TEV protease overnight at 4 °C, followed by addition of Ni-NTA resin (cOmplete His-Tag; Roche) to eliminate TEV protease.

CaM cell pellets were resuspended in Ni-NTA buffer (20 mM HEPES pH 7.5, 200 mM NaCl, 10% v/v glycerol), supplemented with 10 mM imidazole (pH 8.0), EDTA-free cOmplete Protease Inhibitor (Roche) and 0.5 mM Bond-Breaker TCEP and were lysed by sonication, before the lysate was clarified by centrifugation (40,000 × *g*, 45 min, 4 °C). The clarified lysate was incubated with Ni-NTA resin (Roche) pre-equilibrated in Ni-NTA buffer with 5 mM imidazole for >1 h on rollers at 4 °C, before the beads were pelleted at 500 × *g* and washed extensively with Ni-NTA buffer containing 35 mM imidazole. The proteins were eluted in Ni-NTA buffer containing 250 mM imidazole. The eluate was further purified by cleaving the His_6_ tag using recombinant His_6_-TEV protease (for applications where the absence of His_6_ tag was desirable), dialysis, Ni-NTA resin addition to eliminate uncleaved material and the TEV protease. Both cleaved samples were spin concentrated (10 or 30 kDa MWCO; Millipore), before being loaded onto a HiLoad 16/160 Superdex 200 pg column (Cytiva) pre-equilibrated with SEC buffer (20 mM HEPES pH 7.5, 200 mM NaCl, 5% v/v glycerol). Purified fractions, as assessed following resolution by reducing StainFree SDS-PAGE gel electrophoresis (Bio-Rad), were pooled and spin concentrated to 5 mg/mL (CHK2; 30 kDa MWCO) or 50 mg/mL (CaM; 10 kDa MWCO) based on A_280_ and extinction coefficient for each protein, aliquoted, and snap frozen in liquid N_2_ for storage at −80 °C.

Full-length CHK2, harboring a C-terminal AviTag for SPR studies, was expressed and purified from *Sf*21 insect cells, due to toxicity in *E. coli*, using established procedures^40, 41^. Briefly, the bacmid was prepared in DH10MultiBac *E. coli* (ATG Biosynthetics) from a pFastBac GST vector encoding a TEV protease-cleavable GST fusion N-terminal to CHK2-AviTag. *Sf*21 insect cells (Merck) were cultured in Insect-XPRESS (Lonza) media. Each bacmid (1μg) was introduced into 0.9 × 10^6^ *Sf*21 cells by Cellfectin II (Thermo Fisher Scientific) mediated transfection in six-well plates using the Bac-to-Bac protocol (Thermo Fisher Scientific). After 4 days of static incubation at 27 °C in a humidified incubator, the resulting P1 baculovirus was harvested and added to 50 mL *Sf*21 cells at 0.5 × 10^6^ cells/mL density at 4% v/v, which were shaken at 27 °C, 130 rpm. The cell density was monitored daily using a haemocytometer slide and maintained at 0.5–3.0 × 10^6^ cells/mL by diluting with fresh Insect-XPRESS media when necessary, until growth arrest (defined as a cell density less than the twice the cell count 1 day prior). Approximately 24 h after growth arrest was recorded, the P2 baculovirus was harvested by collecting the supernatant after pelleting the cells at 500 × *g* for 5 min. P2 baculovirus was added to 0.5 L *Sf*21 cells cultured at 1–1.5 × 10^6^ cells/mL in 2.8 L Fernbach flasks at 0.2% v/v, and cultured at 27 °C, 90 rpm, until 24 h post growth arrest. Cells were harvested at 500 ×*g* and pellets snap frozen in liquid N_2_ and either thawed immediately for lysis or stored at −80 °C. *Sf*21 cell pellets were lysed and purified by following an analogous protocol to GST-tagged CHK2 expressed in *E. coli*.

### Far-Western Overlay Assay

Full-length recombinant CHK2 were spotted onto PVDF membrane, blocked with 5% w/v skim milk powder in TBST, and probed with recombinant His_6_-tagged CaM (produced in-house) and 500 μM CaCl_2_. CaM was detected using rabbit anti-His_6_-HRP-conjugated antibody (Abcam ab1187) as the secondary (1:5000 dilution). Control experiments, in the presence of 10 mM EGTA, to chelate Ca^2+^ ions, were performed as above. Far-westerns were detected by chemiluminescence using a ChemiDoc Imaging System (Bio-Rad).

### CHK2 Kinase Assays

CHK2 activity was determined by measuring the transfer of radiolabelled phosphate from [γ-^32^P]-ATP to a synthetic peptide substrate (CHKtide; KKKVSRSGLYRSPSMPENLNRPR), synthesised from GenScript (New Jersey, USA). Briefly, purified recombinant CHK2 (Full-length or CHK2^210–531^; 100 pM) were incubated in assay buffer (50 mM HEPES, pH 7.4, 1 mM DTT) containing 200 μM Chktide, 100 μM CaCl_2_, 1 μM recombinant CaM (produced in-house), 200 μM [^32^P]-γ-ATP (Perkin Elmer, MA, USA), 5 mM MgCl_2_ (Sigma) in a standard 30 μL assay for 10 min at 30 °C. Reactions were terminated by spotting 15 μL onto phosphocellulose paper (SVI-P; St Vincent’s Institute, Melbourne) and washing extensively in 1% phosphoric acid (Sigma). Radioactivity was quantified by liquid scintillation counting. CHK2 activity was also measured using an ADP-Glo kinase assay kit (Promega), following standard procedures. Briefly, a 10 μL kinase reaction was assembled, containing 50 ng CHK2, 250 μM CHKtide, 100 μM CaCl_2_, 1 μM recombinant CaM (produced in-house), and 200 μM ATP in kinase buffer (50 mM HEPES pH 7.5, 20 mM MgCl_2_, 2 mM MnCl_2_, 1 mM TCEP). Each reaction was performed at 30 °C for 30 min and terminated by the addition of the ADP-Glo reagent. Following a 40 min with the ADP-Glo reagent, the Kinase Detection reagent was added and luminescence was detected with a microplate reader (CLARIOstar, BMG LabTech) after 30 min.

### Site-specific biotinylation of CHK2-AviTag using recombinant BirA

Site-specific biotinylation of CHK2-AviTag (ASSSSSGLNDIFEAQKIEWHE) was carried out with His-Trx-BirA (produced in *E. coli* in-house), using established procedures^42^. Briefly, CHK2-AviTag was first dialysed into a low salt buffer consisting of 20 mM HEPES pH 7.5, 200 mM NaCl and 1 mM TCEP. CHK2 (100 µM in 952 µL of low salt buffer) was then mixed with magnesium chloride (5 µL of 1M solution; Sigma), adenosine triphosphate (20 µL of 100 mM solution; Sigma), thawed GST-BirA enzyme (20 µL of 50 µM solution) and D-Biotin (3 µL of 50 mM solution in 100 % DMSO; Sigma) and incubated for 1 h minimum at 30 °C in an incubator with gentle shaking (90 rpm). After >1 h, an additional equivalent of His-Trx-BirA and D-biotin were added and the reaction incubated for a further 1 h at 30 °C. Next, 100 µL of a 50 % slurry of Ni-NTA resin (Roche; pre-equilibrated into low salt buffer) was added to the reaction and incubated for 30 min at 4 °C to capture His-Trx-BirA. The resin was then pelleted at 500 ×*g* and the supernatant spin concentrated (30 kDa MWCO; Millipore), before being loaded onto a HiLoad 16/160 Superdex 200 pg column (Cytiva) pre-equilibrated with SEC buffer (20 mM HEPES pH 7.5, 200 mM NaCl, 5% v/v glycerol). The extent of biotinylation was evaluated by mixing with recombinant streptavidin before subjecting to reducing SDS-PAGE with StainFree imaging. With this assay, the gel shift indicated biotinylation was near stoichiometric. Biotinylated full-length CHK2 was concentrated to 2 mg/mL, aliquoted, and snap frozen in liquid N_2_ for storage at −80 °C.

### Surface plasmon resonance

SPR were performed using a BIAcore 8K+ instrument using a SA sensor chip (Cytiva) at 25 °C. Biotinylated full length CHK2 (0.005 mg/mL) was immobilised in a buffer consisting of 20 mM HEPES pH 7.5 and 200 mM NaCl. Direct binding experiments were performed in running buffer (20 mM HEPES pH 7.5, 200 mM NaCl, 2 mM CaCl_2_ and 0.005% Tween) with 240 sec injections and 200 sec dissociations at 25 µL/min, using recombinant CaM (0-200 µM; produced in-house) as the analyte. Flow cells were regenerated in 0.5 M NaCl with 30 sec injections at 30 µL/min. Steady-state responses were determined in affinity evaluation mode using the BIAevaluation software and values plotted as a function of CaM concentration. All affinity measurements were performed in at least two independent experiments using independent protein preparations.

### Thermal Shift Assays

Thermal shift assays were performed as described previously^43^ using a Corbett Real Time PCR machine with proteins diluted in 20 mM HEPES, pH 7.5, 200 mM NaCl to 10 μg in a total reaction volume of 25 μL. SYPRO Orange (Molecular Probes, CA) was used as a probe with fluorescence detected at 530 nm. Two independent assays were performed for wild-type and mutant CHK2 constructs; averaged data are shown for each in **Supp. Fig. 1C**.

### Chemical crosslinking sample preparation

For DMTMM crosslinking, full-length recombinant CHK2 and CaM (1:5 molar ratio) were initially incubated on ice in 50 mM HEPES (pH 7.5), 2 mM CaCl_2_ for 30 min. The protein mixture (∼3 μg) was then incubated for 30 min at room temperature with a range of DMTMM concentrations (0, 5, 10, 20, 50 and 100 mM; Sigma, # 74104), which was prepared in 50 mM HEPES (pH 7.5). Following incubation, each reaction was quenched with 100 mM Tris-HCl (pH 7.5), mixed with reducing sample buffer, heated to 100°C for 5 min, then resolved by reducing SDS-PAGE gel electrophoresis (Bio-Rad). SDS-PAGE gels were stained using SimplyBlue SafeStain (Thermo Fisher). For SDA crosslinking, recombinant full-length CHK2 and CaM (1:2 molar ratio) were initially incubated on ice in 50 mM HEPES (pH 7.5), 2 mM CaCl_2_ for 30 min. Protein (1.5 mg/mL) was then mixed with SDA (NHS-Diazirine; 1 mg/mL; Thermo Fisher, # 26167) and incubated in the dark for 30 min at room temperature to react the NHS-ester group. The diazarine group was then photo-activated using ultraviolet light irradiation (UVP CL-1000L UV cross-linker) at 365 nm. Samples were added to upturned lids excised from 1.5 mL microfuge tubes and placed on ice at a distance of 5 cm from the lamp and irradiated for 1 min. The reaction mixtures from the titration were combined and quenched with 100 mM Tris-HCl (pH 7.5), mixed with reducing sample buffer, heated to 100°C for 5 min, then resolved by reducing SDS-PAGE gel electrophoresis (Bio-Rad). SDS-PAGE gels were stained using SimplyBlue SafeStain (Thermo Fisher) and crosslinked adducts excised for mass spectrometry analysis.

### Chemical crosslinking sample preparation for mass spectrometry-based proteomics

Protein gel bands were excised (based on their migration relative to the MW marker) and destained using 50% acetonitrile for 30 min and 37 °C. The gel band was then dehydrated by adding 100% acetonitrile (ACN) and incubating at room temperature for 10 min, before aspirating the ACN and further drying gel piece with the vacuum centrifuge (CentriVap, Labconco). Proteins were reduced by the addition of 1 mM dithiothreitol (DTT) in 50 mM ammonium bicarbonate for 30 min. Excess DTT was aspirated before the addition of 55 mM iodoacetamide to alkylate the sample for 60 min at room temperature. The gel slice was then washed with 50% acetonitrile twice and 100% ACN once before drying to completion in the vacuum centrifuge. Proteins were digested overnight with 500 ng trypsin in 50 mM ammonium bicarbonate at 37 °C, and extracted the following day using 60% acetonitrile/0.1% formic acid. The collected peptides were lyophilized to dryness using a CentriVap (Labconco), before reconstituting in 10 µL 0.1% formic acid/2% ACN ready for mass spectrometry analysis.

### Chemical crosslinking mass spectrometry analysis

Reconstituted peptides were analyzed on Orbitrap Eclipse Tribrid mass spectrometer interfaced with Neo Vanquish liquid chromatography system. Samples were loaded onto a C18 fused silica column (inner diameter 75 µm, OD 360 µm × 15 cm length, 1.6 µm C18 beads) packed into an emitter tip (IonOpticks) using pressure-controlled loading with a maximum pressure of 1,500 bar using Easy nLC source and electro sprayed directly into the mass spectrometer.

We first employed a linear gradient of 3-30% of solvent-B at 400 nL/min flow rate (solvent-B: 99.9% (v/v) ACN) for 100 min, followed by a gradient of 30-40% solvent-B for 20 min and 35-99% solvent-B for 5 min. The column was then maintained at 99% B for 10 min before being washed with 3% solvent-B for another 10 min comprising a total of 145 min run with a 120 min gradient in a data dependent (DDA) mode. MS1 spectra were acquired in the Orbitrap (R = 120k; normalised AGC target = standard; MaxIT = Auto; RF Lens = 30%; scan range = 380–1400; profile data). Dynamic exclusion was employed for 30 s excluding all charge states for a given precursor. Data dependent MS2 spectra were collected in the Orbitrap for precursors with charge states 3-8 (R = 50k; HCD collision energy mode = assisted; normalized HCD collision energies = 25%, 30%; scan range mode = normal; normalised AGC target = 200%; MaxIT = 150 ms).

### Chemical crosslinking raw data processing and peptide identification

Raw data files were converted to MGF files using MS convert^44^. For DMTMM crosslinking analysis, peak files were searched using the MeroX software^45^ (version 2.0.1.4) and a fasta file containing the full-length human CHK2 and CaM sequences. K to D or E residues were set as specificity sites for DMTMM crosslinking (-H_2_O). Trypsin was set as the enzyme allowing for three missed cleavages. Maximum peptide length was set to 40. Precursor precision was set at 10 ppm with fragment ion precision set at 20 ppm. For SDA crosslinking, MGF files were searched against a fasta file containing the CHK2 and CaM sequences sequence using XiSearch software^46^ (version 1.7.6.7) with the following settings: crosslinker = multiple, SDA and noncovalent; fixed modifications = Carbamidomethylation (C); variable modifications = oxidation (M), SDA-loop (KSTY) DELTAMASS:82.04186484, SDA-hydro (KSTY) DELTAMASS:100.052430; MS1 tolerance = 6.0ppm, MS2 tolerance = 20.0ppm; losses = H_2_O,NH_3_, CH_3_SOH, CleavableCrossLinkerPeptide:MASS:82.04186484). FDR was performed with the in-built xiFDR set to 5%. Data were visualized using the XiView software^47^.

### Computational Modelling of CHK2 and CaM

The sequence of full-length human CHK2 (Uniprot O96017) and full-length human CaM (Uniprot P0DP23) were used to generate a complex model using ColabFold v1.5.5: AlphaFold2 using MMseqs2^48^.

### Hydrogen-deuterium exchange (HDX) mass spectrometry sample preparation

HDX reactions comparing apo CHK2 to CHK2 incubated with CaM were carried out in a 10 µl reaction volume containing 10 pmol of full-length human CHK2 and 150 pmol of human CaM. Both proteins were recombinantly expressed (described above). The exchange reactions were initiated by the addition of 6.0 µL of D_2_O buffer (20 mM HEPES pH 7.5, 200 mM NaCl, 2mM CaCl_2_, 92.4% D_2_O (v/v)) to 4.0 µL of protein (final D_2_O concentration of 55.5%). Reactions proceeded for 3 s, 30 s, 300 s and 3000 s at 20 °C before being quenched with ice-cold acidic quench buffer, resulting in a final concentration of 0.6M guanidine HCl and 0.9% formic acid post quench. All conditions and timepoints were created and run in independent triplicate. Samples were flash frozen immediately after quenching and stored at −80 °C.

### HDX protein digestion and MS/MS data collection

Protein samples were rapidly thawed and injected onto an integrated fluidics system containing a HDx-3 PAL liquid handling robot and climate-controlled (2 °C) chromatography system (LEAP Technologies), a Waters ACQUITY UPLC I-Class Series System, as well as an Impact HD QTOF Mass spectrometer (Bruker) using an established procedure^49^. The samples were run over an immobilized pepsin column (Affipro; AP-PC-001) at 200 µL/min for 4 min at 2 °C. The resulting peptides were collected and desalted on a C18 trap column (ACQUITY UPLC BEH C18 1.7 µm column, 2.1 mm × 5 mm; Waters 186004629). The trap was subsequently eluted in line with an ACQUITY 300Å, 1.7 µm particle, 100 mm × 2.1 mm C18 UPLC column (Waters; 186003686), using a gradient of 3-35% Buffer (Buffer A 0.1% formic acid; Buffer B 100% acetonitrile) over 11 min immediately followed by a gradient of 35-80% over 5 min. Mass spectrometry experiments acquired over a mass range from 150 to 2200 m/z using an electrospray ionization source operated at a temperature of 200 °C and a spray voltage of 4.5 kV.

### HDX peptide identification

Peptides were identified from the non-deuterated samples of CHK2 using data-dependent acquisition following tandem MS/MS experiments (0.5 s precursor scan from 150-2000 m/z; twelve 0.25 s fragment scans from 150-2000 m/z). MS/MS datasets were analysed using FragPipe v19.1 and peptide identification was carried out by using a false discovery-based approach using a database of purified proteins and known contaminants^50, 51^. MSFragger was utilized^52^, and the precursor mass tolerance error was set to −20 to 20ppm. The fragment mass tolerance was set at 20ppm. Protein digestion was set as nonspecific, searching between lengths of 4 and 50 aa, with a mass range of 400 to 5000 Da.

### HDX mass analysis of peptide centroids and measurement of deuterium incorporation

HD-Examiner Software (Sierra Analytics) was used to automatically calculate the level of deuterium incorporation into each peptide. All peptides were manually inspected for correct charge state, correct retention time, appropriate selection of isotopic distribution, etc. Deuteration levels were calculated using the centroid of the experimental isotope clusters. Results are presented as relative levels of deuterium incorporation and the only control for back exchange was the level of deuterium present in the buffer (55.5%). Differences in exchange in a peptide were considered significant if they met all three of the following criteria: ≥5% change in exchange, ≥0.40 Da difference in exchange, and a p value <0.01 using a two tailed student t-test. Samples were only compared within a single experiment and were never compared to experiments completed at a different time with a different final D_2_O level. The data analysis statistics for all HDX-MS experiments are in **Supp. Table 2** according to published guidelines^53^. The mass spectrometry proteomics data have been deposited to the ProteomeXchange Consortium via the PRIDE partner repository^54^ with the dataset identifier (PXD048473).

### Bioinformatic analysis

CHK2 homolog sequences were gathered using BlastP against the NRAA database, aligned with ClustalOmega^55^ and visualized with Jalview^56^

### RPE cell culture

hTERT-immortalized RPE cells hTERT RPE-1 (ATCC, #CRL-4000) cells were maintained in DMEM/F12 (1:1) (Gibco, #11330-032) supplemented with 10% FBS (Bovogen, #SFBS-F) and 0.01 mg/ml hygromycin B (Roche, #10843555001) in an incubator at 37°C and 5% CO2. Cells were passaged twice weekly.

### Generation and screening of CRISPR-edited cell lines

hTERT RPE-1 cells were edited using CRISPR editing strategy described previously^57^. CHK2 guide RNA sequences were designed using the CRISPOR software package^58^. 20 bp guides beginning with G were selected. 5′-CACC-3′ and 5′-AAAC-3′ were appended to the sense and antisense guide sequences, respectively. Annealed gRNAs were cloned into the eSpCas9-hGeminin plasmid (Addgene plasmid, #86613). The resulting plasmid is referred to as eSpCas9-hGeminin –CHK2 gRNA. To generate the CHK2 K373A point mutation a 200 bp single-stranded oligonucleotide (ssODN) donors was synthesised by IDT, which included lysine 373 to alanine mutation, silent mutations in the PAM (generates an AclI restriction site used for screening purposes) and in the corresponding gRNA sequence (to reduce re-editing).

For transfection, cells were harvested by trypsinization at 80% confluence. 1 × 10^6^ cells were co-transfected with 5 µg of eSpCas9-hGeminin –CHK2 gRNA plasmid, 5 µg of ATP1A1_plasmid_donor_RD (Addgene plasmid, #86551), and 2.5 µl ssODN donor (100 µM). To generate control cell lines, 1 × 10^6^ cells were co-transfected with 5 µg of eSpCas9-hGeminin plasmid (Addgene plasmid, #86613) and 5 ug of ATP1A1_plasmid_donor_RD (Addgene plasmid, #86551). Cells were electroporated using the Neon Transfection System (Invitrogen, #MPK5000) at 1350 V, 20 ms, 2 pulse. Electroporated cells were incubated in 15 cm round dishes for 72 h, then treated with 10 µM ouabain (sigma, #O3125) selection media, which was replaced every few days. To obtain single clones including the point mutations, following 7 days of ouabain treatment, cells were trypsinization and limited dilutions were performed. Colonies in 96 wells were expanded to six-well plates for cryopreservation and DNA analysis.

Genomic DNA were isolated by using Genomic DNA purification kit (Sangon, #B618503). To screen cell lines for genome edits corresponding to CHK2 lysine 373, a 684 bp PCR product was amplified using the primer Forward: 5′-GAGACACTGGGGTCTAAGAACCATGTAG −3′ and Reverse: 5′-GGTGGTGTGCATCTGTAGTCCCAG −3′. The PCR product was digested with AclI (New England BioLabs, #R0598S) to confirm insertion of the ssODN. To validate the point mutation, PCR products were sequenced using the forward primer 5′-GAGACACTGGGGTCTAAGAACCATGTAG −3′ by Sanger Sequencing

### Cell proliferation assays

For cell growth assays, hTERT-immortalised cells were plated at 2000 cells per well (control cells) and 1000 cells per well (*CHK2^K373A^*) in a 96-well plate and left to settle for 24 h before being transferred to an IncuCyte S3 instrument. Images were taken every 2 h for 72 h by phase contrast. Cell proliferation was then measured using IncuCyte S3 software (version 2022B). To examine differences in the presence of DNA damage, cells were stimulated at time 0 with Camptothecin (CPT; Sigma) at 10 and 100nM prepared in DMSO. Cell confluency was measured as described above. GraphPad Prism 7 was used for data analysis. Paired t-test or two-way ANOVA were used for calculating significance.

## DATA AVAILABILITY

Source data are published as an accompanying file; any additional data, including expression construct sequences, are available from the corresponding authors upon request. Any materials are available from the corresponding authors under Materials Transfer Agreement. The mass spectrometry proteomics data have been deposited to the ProteomeXchange Consortium via the PRIDE partner repository^54^ with the dataset identifier (PXD048473).

## ACKNOWLEGDMENTS

We thank James Knox and the MiMo crew for facilitating vital discussions, and Meredith Jenkins for assistance with the HDX-MS experiments. We thank the National Health and Medical Research Council of Australia for grant (JMM, 1172929; JWS, 2001817) and infrastructure (9000719) support, the Australian Research Council for grant support (JWS, DP210102840) and the Victorian State Government Operational Infrastructure Support Scheme. We thank the Cancer Research Society (Operating Grant 1052949) and the Michael Smith Foundation for Health Research (Scholar award 17868) for support to JEB.

## AUTHOR CONTRIBUTIONS

C.R.H. designed, performed, and analysed experiments; C.R.H. wrote the paper with J.M.M.; T.W. and J.P. generated the CRISPR-edited cell lines; S.N.Y., T.D., H.N., S.S., K.A.D., L.M., L.F.D., G.M., A.R.M., and J.E.B. performed and analysed experiments; J.W.S. and J.M.M. supervised the project, analysed, and interpreted data; all authors commented on the manuscript.

