## Supplemental Tables 1-2; Supplemental Figures 1-3 for "Unconventional binding of Calmodulin to CHK2 kinase inhibits catalytic activity"

### SUPPLEMENTARY TABLES

#### Supplementary Table 1. Oligonucleotide sequences

| | |
|---|---|
| CHK2 2 IF fwd | 5’-GTATTTTCAGGGATCCTCTCGGGAGTCGGATGTTGAG-3’ |
| CHK2 553 STOP IF rev | 5’-CAGTCACGATGAATTCACAACACAGCAGCACACACAG-3’ |
| CHK2 210 IF fwd | 5’-GTATTTTCAGGGATCCTCAGTTTATCCTAAGGCATTAAGAG-3’ |
| CHK2 531 STOP IF rev | 5’-CAGTCACGATGAATTCACTCGGCACCCTCGGCTTC-3’ |
| pcDNA XhoI N-FLAG[‡] | 5’-cgcCTCGAGATGGATTACAAGGATGACG-3’ |
| pcDNA STOP HindIII rev | 5’-cgcAAGCTTTCACAACACAGCAGCAC-3’ |
| pFB CHK2 IF fwd | 5’-ACCATGTCGTACTACatgtcccctatactaggttattggaaaattaagg-3’ |
| pFB CHK2 IF rev | 5’-GATGTCGTTCAGACCTCACAACACAGCAGCACACACA-3’ |
| pFB vector fwd | 5’-GGTCTGAACGACATCTTCGAGG-3’ |
| pFB vector rev | 5’-GTAGTACGACATGGTTTCGGACCG-3’ |
| hCaM BamHI 2 fwd | 5’-cgcGGATCCgctgatcagctgaccgaagaacag-3’ |
| hCaM 149 STOP NotI rev | 5’-cgcGCGGCCGCTCattttgcagtcatcatctgtacgaattc-3’ |
| CHK2 K373 sense | 5’-aaacTATGTGGAACCCCCACCTAC-3’ |
| CHK2 K373 antisense | 5’-caccGTAGGTGGGGGTTCCACATA-3’ |
| CHK2 CRISPR screen fwd | 5′-GAGACACTGGGGTCTAAGAACCATGTAG-3′ |
| CHK2 CRISPR screen rev | 5′-GGTGGTGTGCATCTGTAGTCCCAG-3′ |
| CRISPR sequence fwd | 5′-GAGACACTGGGGTCTAAGAACCATGTAG-3′ |

[‡] Restriction sites underlined

**Supplementary Table 2. HDX-MS data analysis statistics**

| | CHK2 apo | CHK2 + CaM (15μM) |
|---|---|---|
| HDX reaction details | %$D_2O$ = 55.5%<br>$pH_{(read)}$ = 7.5<br>Temperature = 20 °C | %$D_2O$ = 55.5%<br>$pH_{(read)}$ = 7.5<br>Temperature = 20 °C |
| HDX time course | 3s at 0C, 3s, 30s, 300s, 3000s at 18C | 3s at 0C, 3s, 30s, 300s, 3000s at 18C |
| HDX controls | N/A | N/A |
| Back-exchange | Corrected based off %$D_2O$ | Corrected based off %$D_2O$ |
| Number of unique peptides | 85 | 85 |
| Sequence coverage | 77.4% | 77.4% |
| Average peptide length | 14.1 | 14.1 |
| Average peptide redundancy | 2.2 | 2.2 |
| Replicates | 3 | 3 |
| Repeatability | Average StDev = 0.8% | Average StDev = 0.8% |
| Significant differences in HDX | >5% and >0.4 Da and unpaired t-test <0.01 | >5% and >0.4 Da and unpaired t-test <0.01 |

### SUPPLEMENTARY FIGURES

**Supplementary Figure 1 | Activity, purification, and stability of recombinant CHK2 constructs. (A)** In vitro kinase activities of wild-type CHK2 (full-length and 210-531), measured by ADP-Glo assay, in presence or absence of $Ca^{2+}$-CaM. Individual data points are plotted; the bar and error bars shown represent mean ± SD of three independent assays. Statistical analysis was performed by two-way ANOVA; **** signifies P<0.0001. **(B)** Reducing SDS-PAGE analysis of the purified CHK2 full length and kinase domain (210-531) constructs visualized by stain-free imaging. Positions of molecular weight standards (first lane) are annotated on the left of the gel. The purity of the shown protein preparations are representative of at least 3 independent expressions and purifications for each construct. **(C)** Thermal shift curves of purified CHK2 full length and kinase domain (amino acids 210-531) constructs, performed using differential scanning fluorimetry confirms that these variants are folded. Data represent mean ± SD of at least two independent experiments. Data are plotted throughout for wild-type CHK2 (full length) in black, wild-type CHK2 (210-531) in pink, and mutants are color-coded: K245A in purple; K255A, light blue; K373A, dark blue; K437A, green; and K465A, brown. K249A (kinase dead) is not shown; high background fluorescence prevented $T_m$ estimation, and suggests partial unfolding.

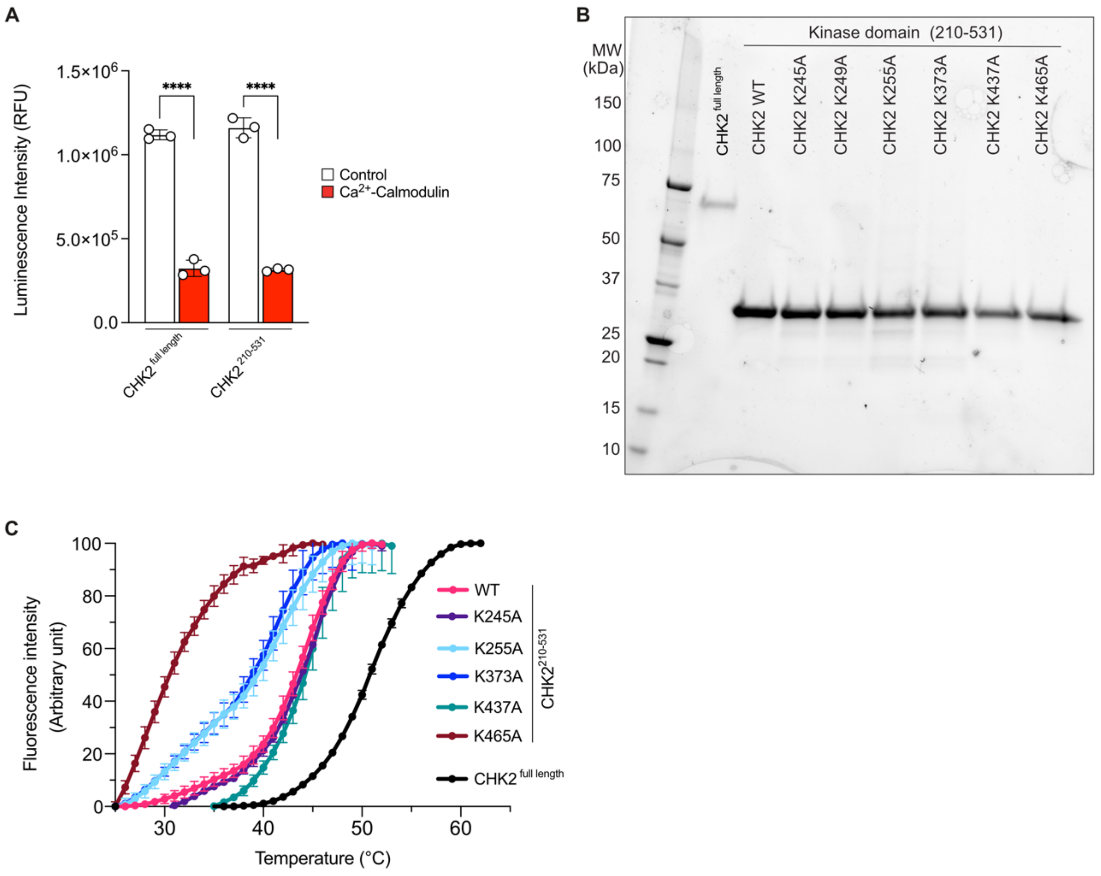

**Supplementary Figure 2 | Direct interaction between CHK2 and $Ca^{2+}$-CaM. (A-B)** Double referenced SPR sensorgrams for full length CHK2. Two independent repeats are shown, where the sensorgram is color-coded based on concentration. **(C-D)** Steady state analysis for (A; blue) and (B; red) SPR experiments are shown, along with the dissociation constant.

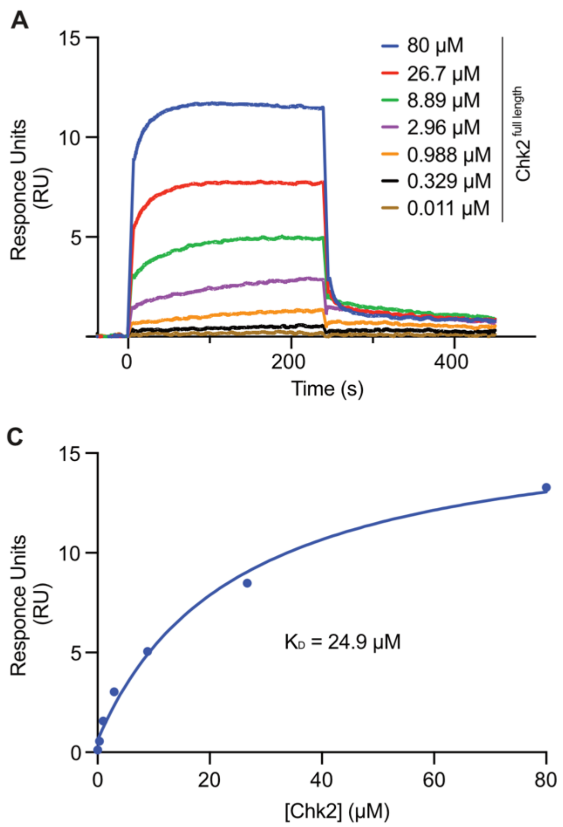

**Supplementary Figure 3 | HDX-MS deuterium uptake plots.** Representative CHK2 peptides (black) displaying decreases in deuterium exchange upon binding Calmodulin (red). Data are presented as mean values ± SD from three independent experiments (n=3). Most error bars are smaller than the size of the point. Source data are provided as a Source Data file.

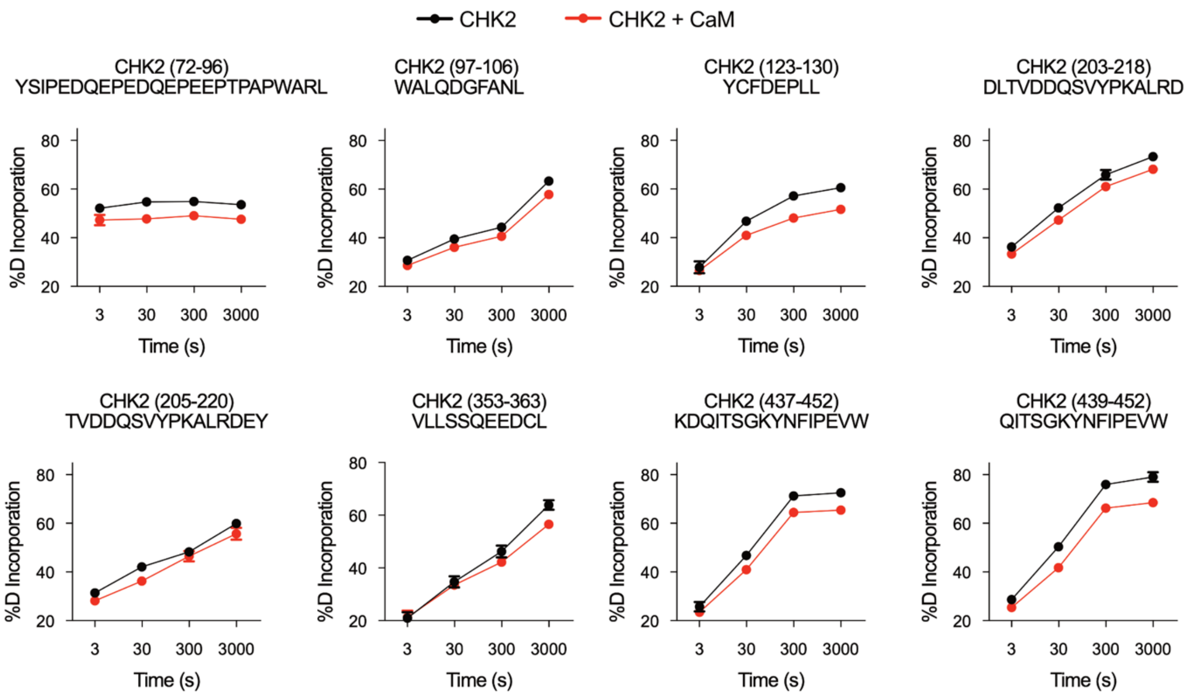

**Supplementary Figure 4 | Genotypic validation of CRISPR-edited RPE cells.** The endogenous *CHK2* locus of hTERT immortalized RPE cells was CRISPR edited using the lower donor sequence encoding the K373A mutation (blue) and silent substitutions (green) around the PAM site (yellow). Figure generated using Biorender.

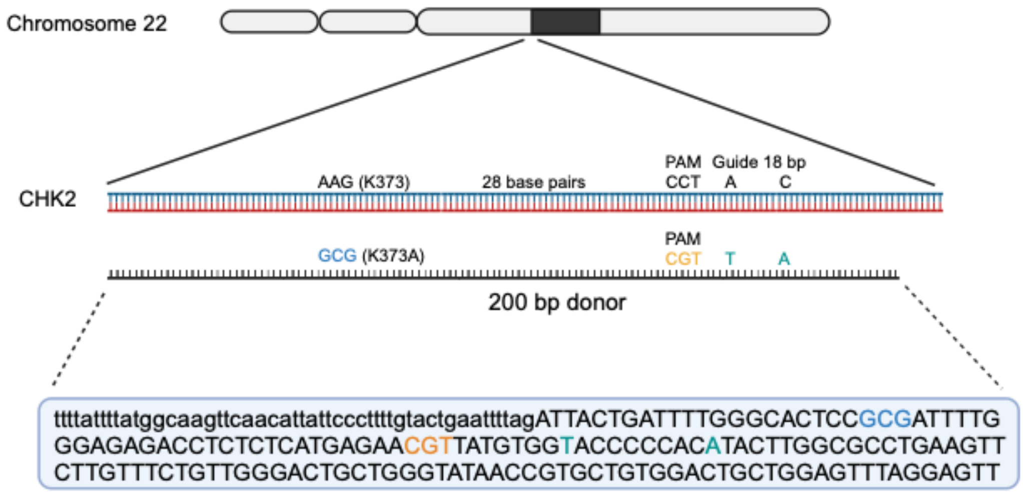
